# Universality of evolutionary trajectories under arbitrary competition dynamics

**DOI:** 10.1101/2021.06.17.448795

**Authors:** Andrea Mazzolini, Jacopo Grilli

## Abstract

The assumption of constant population size is central in population genetics. It led to a large body of results, that are robust to modeling choices and that have proven successful to understand evolutionary dynamics. In reality, allele frequencies and population size are both determined by the interaction between a population and the environment. Relaxing the constant-population assumption have two big drawbacks. It increases the technical difficulty of the analysis, and it requires specifying a mechanism for the saturation of the population size, possibly making the results contingent on model details. Here, we develop a framework that encompasses a great variety of systems with an arbitrary mechanism for population growth limitation. By using techniques based on scale separation for stochastic processes, we are able to calculate analytically properties of evolutionary trajectories, such as the fixation probability. Remarkably, these properties assume a universal form with respect to our framework, which depends on only three parameters related to the inter-generation timescale, the invasion fitness, and the carrying capacity of the strains. In other words, different systems, such as Lotka-Volterra or a chemostat model (contained in our framework), share the same evolutionary outcomes after a proper re-mapping of their parameters. An important and surprising consequence of our results is that the direction of selection can be inverted, with a population evolving to reach lower values of invasion fitness.

## I. INTRODUCTION

Competition is a fundamental agent of natural selection. The scarcity of resources (nutrient, water, space, …) limits population growth, determining a selection pressure. Variants — strains in the following — that grow faster or more efficiently spread within the population. The long history of mathematical population genetics has produced several seminal results that quantitatively describe these processes. The vast majority of results are based on the assumption of constant population size [1–8]. The success of this assumption lies in the generality of the results. When the population size is large enough, the diffusion limit developed independently by Wright and Kolmogorov, pioneered in population genetics by Malécot[63] and Kimura [1, 9], is the convergence point of several alternative models. In fact, while different population genetics models (Wrigh-Fisher [10, 11], Moran [12], conditional branching processes [13], and some Canning processes [14]) start from radically different assumptions about the genealogical and demographic structure of the population, they share the same predictions, up to a simple rescaling of timescales and parameters. For most of the theoretical advances in population genetics, the total population size is treated as an effective parameter of the model, which should be fitted from data, and the strain frequencies are the only dynamical degrees of freedom. However, mechanistically, not only the strain frequencies but also the total population size is determined by the interaction between a population and the environment. Moreover, experimental works [15, 16] suggest that the variation of the population size can play a role in the evolutionary process, implying that its dynamics should not be neglected in theoretical descriptions.

Non-stationary conditions are an important example where the dynamics of population size cannot be neglected to understand how evolution — here intended as the dynamics of relative frequencies of strains — unfolds. For instance, range expansions leave strong signature on the genetic diversity of a population [17–19].

A classical deterministic approach to couple the population growth and the frequencies of the strains is density-dependent selection [20–23]. Another approach to study evolution under varying total population sizes is to consider the dynamics of the latter as decoupled from the strain frequency, by assuming a priori the dynamics of the total population [24–27]. Here we instead focus on stochastic scenarios. This approaches require to model how environmental constraints limit population growth, for instance by considering a Lotka-Volterra dynamics [28–34] or generalizations of the Moran model [35]. In these kinds of models, the population size is a stochastic variable itself that can fluctuate. Those fluctuations are not independent of strain frequencies, leading to highly non-trivial evolutionary dynamics.

The increased realism of these models undermines, at least in principle, the generality of the standard population genetics results. To what extent do the details of the competition structure determine the evolutionary outcome? For instance, will a population whose growth is described by a logistic model differ in its evolutionary trajectory from a population described by Gompertz growth?

In this work, we address this question by introducing and considering a general ecological framework to describe the dynamics of a haploid asexual population. This approach encompasses several alternative models of competition — from Lotka-Volterra to consumer-resource models in a chemostat, to a Gompertzian growth — under the same mathematical set-up. In other words, this ecological framework defines the mathematical details of how the growth of a population is limited by competition for limited resources.

Given the generality of our framework, one could in principle expect that the evolutionary dynamics depend on the specific details of the model considered to describe population self-limitation. We show that evolutionary predictions are instead robust and insensitive to the details of population dynamics. Within our framework, evolutionary observables, such as the fixation probability and the fixation time, are universal and depend on only three parameters, related to the concepts of inter-generation timescale, invasion fitness, and carrying capacity.

We first present the general framework in the model section. We start to illustrate the deterministic limit, where basic ideas of the framework can be easily grasped. We then derive the stochastic dynamics, described as a system of Langevin equations. In the limit of small fitness differences, a time-scale separation allows us to calculate analytically the properties of the evolutionary dynamics. We describe the behavior of these properties over evolutionary timescales. We also illustrate an explicit example of the chemostat model for resource competition, showing how the previous general results can lead to an evolutionary trajectory with decreasing fitness over time.

## II. MODEL

To help the reader to build its intuition on the presented framework, we first introduce our framework in the case of large population sizes (which allow neglecting stochasticity), in presence of a single clonal population. We then consider the case of finite population size and introduce the stochastic version for a clonal population. We then generalize our framework to multiple strains in the deterministic limit and, finally, multiple strains in the stochastic formulation, which is the framework considered for the rest of the paper.

### A. A general model for the growth of a clonal population in presence of limiting factors

We consider a clonal population whose growth is limited by some resources. The growth curve of a clonal population displays a typical sigmoid shape, as depicted in Fig. 1: a rapid initial growth followed by a deceleration due to nutrient limitation, and, eventually, a convergence of the abundance to a carrying capacity, determined by the availability and quality of nutrients. Multiple models — such as Logistic, Gompertz, and Von Bertalanffy — capture this phenomenology, by explicitly quantifying the dependence of per-capita growth rates on population abundance. These models differ in the specific functional form of the population growth trajectory. For instance, in the Logistic model, the per-capita growth rate decreases linearly with population abundance, while in the Gompertz model it decreases logarithmically.

**FIG. 1:**
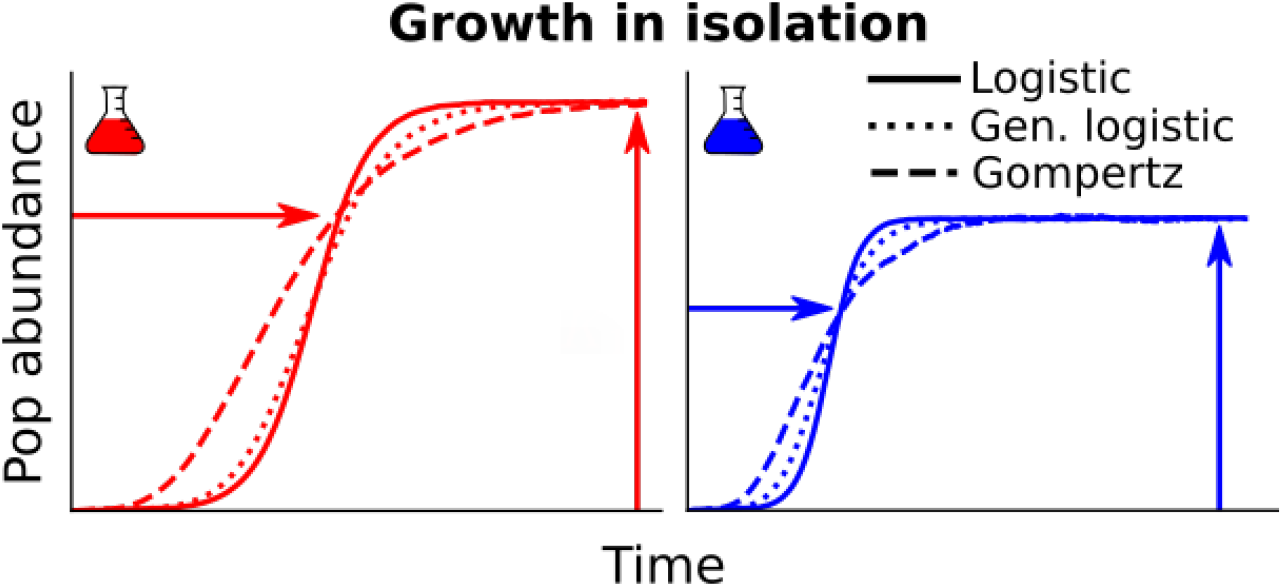
Typical growth curve of a clonal population in isolation, characterized by a carrying capacity *K*_*i*_ and a timescale to reach the saturation *T*_*i*_.

This large class of models can be captured, in full generality, by making the per-capita growth rate depend on the total population abundance *N* through generic functions:

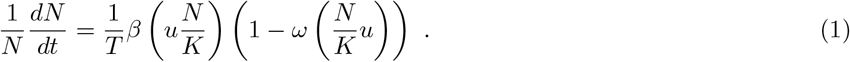

The function *β*(·) and function *ω*(·) determine the specific shape of growth curves, which distinguish between alternative models (see below). The value *u* simply equals to *ω*^*−*1^(1), so that *ω*(*uN/K*) = 1 for *N* = *K*. The two parameters *T* and *K* capture instead the dimensional components of these growth curves. The *carrying capacity K* defines the values of population abundance reached at large times. A *timescale T* sets the speed of convergence of population curves to carrying capacity. Fig. 1 shows the example of two populations with different strains, each characterized by different values of *K* and *T*.

Eq. 1 reduces to standard models under specific choices of the two functions *β*(·) and *ω*(·). Logistic growth is obtained using *β*(*z*) = 1 and *ω*(*z*) = *z*, the chemostat model [36] with *β*(*z*) = 1*/z* and *ω*(*z*) = *z*, Gompertz growth for *β*(*z*) = 1 and *ω*(*z*) = log(*z*), and Von Bertalanffy model for *β*(*z*) = *z*^*α−*1^ and *ω*(*z*) = *z*^1*−α*^ (see Appendix A).

To enforce the saturating phenomenology shown in Fig. 1, few constraints are required for the two generic functions. Specifically, that both are positive and that *ω*(·) is monotonically increasing. This choice guarantees the existence of a unique, globally stable, fixed point *N* ^*∗*^ = *K*, to which the population abundance converges for large times. These choices can be relaxed in presence of more complex biological and ecological mechanisms affecting population growth, that we will not consider in this manuscript. For instance, in presence of a strong Allee effect [37], the per-capita growth rate would be negative at small population abundance, corresponding to a non-monotonicity of *ω*(·).

### B. A general birth-death model for the growth of a clonal population

Eq. 1 can be seen as the continuous deterministic limit, obtained for large population sizes, of a discrete stochastic process that describes the dynamics of abundance of a finite number of individuals. It is natural to define such a model as a birth-death process with microscopic rates

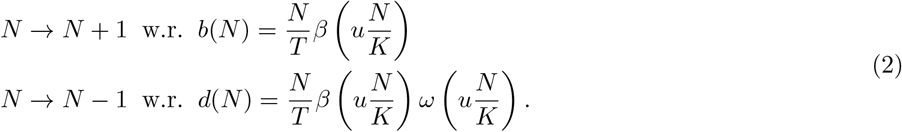

This formulation further clarifies the interpretation of the parameters and the functions appearing in eq. 1. The value of *T* sets the unique timescale of the process, to which both the birth and death rate are inversely proportional, and is directly related to the generation time (see appendix C for more details). The function *β*(·) describes the density dependence of the birth rate, while the function *ω*(·) sets the dependence of the ratio between birth and death rate. The constraints on the two functions that have been introduced in the deterministic case have to be valid also in this context. First, *β*(·) and *ω*(·) must be positive since the rates must be positive. Second, the monotonicity of *ω*(·) enforces the population saturation: the death rate exceed the birth rate when 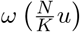 becomes larger than 1, i.e., for *N > K*. The phenomenology of this class of stochastic models have been studied for specific choices of *β*(·) and *ω*(·). In particular, extensive work has been done for logistic growth (*β*(*z*) = 1 and *ω*(*z*) = *z*) [38].

The stochastic dynamics of *N* can be divided into three temporal phases. The first phase is dominated by a transient, which depends on the initial condition and directly corresponds to the transient of the deterministic dynamics depicted in Fig. 1. In the case of logistic growth, this first phase requires a time of the scale *T* log(*K*). The second phase corresponds to fluctuations around *N* = *K* and displays a variability of the order of 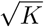 *K*. Approximation for the quasi-stationary distribution of these fluctuations is well known [39, 40]. The timescale *T* determines the typical auto-correlation time of these fluctuations. The stability of the second phase is only apparent, as it is a meta-stable phase. The stochastic process defined in eq. 2 has an absorbing state in *N* = 0 and very rare fluctuations can drive the population to extinction, corresponding to the third phase. The rate of extinction is very small for large populations. The typical time to extinction — which, according to our definitions, corresponds to the duration of the second phase — scales exponentially with *K*. In the case of the logistic model [38] this time is already astronomical for relatively small population sizes. For instance, for *K* = 100 the average time to extinction is on the order of 10^32^ generation times.

### C. Deterministic limit in the case of multiple strains

We want now to focus on the evolutionary dynamics of different strains that compete with each other, by studying how intra-population variability changes over time. We consider therefore *A* strains that differ in their growth rates through their demographic parameters. While *K* and *T* are enough to specify the scales of the growth curves obtained in isolation, they are in fact generally not enough to describe the dynamics in co-cultures. Fig. 2 depicts the setting of a pairwise competition experiment, where a single individual (or a small population) of a mutant is inoculated in a large resident population. In absence of multiple substitutable resources (or other factors giving rise to frequency-dependent selection), only one of the two strains — the one with the higher fitness — survives.

**FIG. 2:**
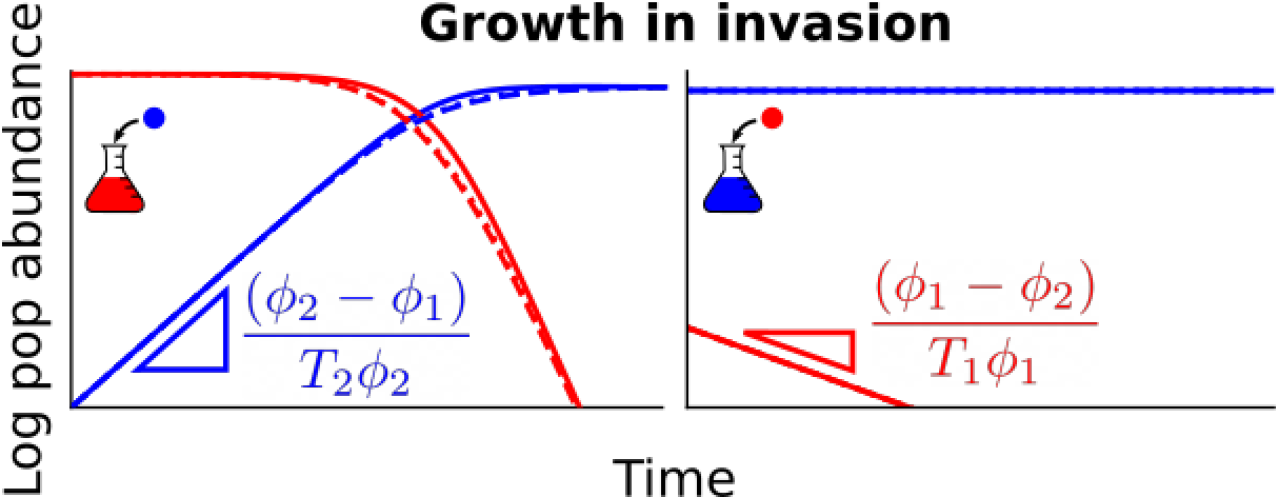
Typical growth curves of two competitive strains, where one tries to invade tho other one at carrying capacity. The invasion success is determined by the invasion fitness *ϕ*_*i*_. The lines are simulated with equation 3, for different choices of the generic functions as in figure 1.

To include this feature in our model, the deterministic equation for the abundance of strain *i* reads

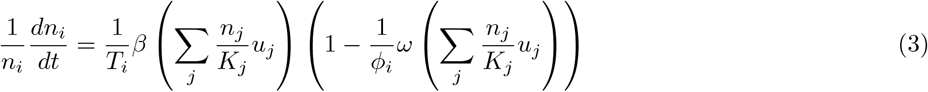

where *u*_*i*_ = *ω*^*−*1^(*ϕ*_*i*_). The generic functions *β*(·) and *ω*(·) are independent of the strain and obey the same constraints as before. As described above, the values of *T*_*i*_ and *K*_*i*_ define the growth curve of that particular strain in a clonal population as shown in Fig. 1. The new crucial ingredient here is the parameter *ϕ*_*i*_, which is related to the (invasion) fitness of strain *i*.

As shown in Appendix B, this system of non-linear differential equations admits a unique, globally stable, fixed point, where only one strain survives reaching a population abundance equal to its carrying capacity. The strain that survives is the one characterized by the largest value of *ϕ*, which we, therefore, identify as the fitness of a strain.

Consistently with the existence of a globally stable equilibrium, the outcomes of invasion experiments depend only on the fitness *ϕ* of the strains involved. In particular, the growth rate of the mutant strain (usually called “invasion fitness”) is proportional to the difference between its fitness *ϕ*, and the one of the resident population (see Figure 2 and Appendix C). It is important to remark that, in general, the invasion fitness does not correspond to the Malthusian fitness [41] (i.e. the per-capita growth rate in exponential growth) as the latter refers only to growth in isolation. For instance, the Malthusian fitness in the logistic model is equal to 1*/T*_*i*_. Therefore the strain that is expected to out-compete the others (the one with the highest value of *ϕ*) is not necessarily the one with the higher growth rate in the exponential phase (1*/T* for the logistic model).

A particularly simple and paradigmatic case occurs when *T*_*i*_ = 1*/*(*τϕ*_*i*_) [64] and *K*_*i*_ = *K* for all *i*. In that case, in fact, the dynamics of the total population abundance has a globally stable fixed point *N* ^*∗*^ = *K* independently of the relative abundances of individual strains. This allows to write explicitly the dynamics for the relative population abundances *x*_*i*_ = *n*_*i*_*/N*, which, by imposing *N* = *K*, turns out to correspond to the replicator equation

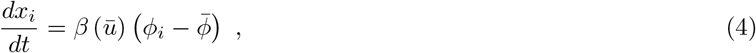

where 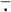 denotes the average over the population (e.g. 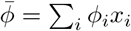).

### D. Birth-death model for multiple strains

Finally, we generalize therefore eq. 2 to describe the coupled birth-death process of multiple strains, whose deterministic dynamics is eq. 3. The birth and death rates *b*_*i*_(**n**) and *d*_*i*_(**n**) depend on the abundance of all the other strains, **n** = *n*_1_, …, *n*_*A*_, as all the individual compete — even if with different ability — for the same limited resources. The rates read

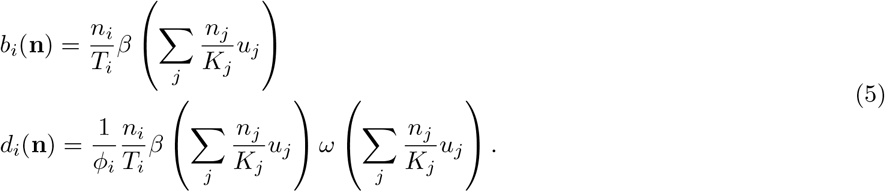

If *ϕ*_*i*_ = 1, *T*_*i*_ = *T*, and *K*_*i*_ = *K* for all *i*, the population abundance *N* = ∑_*j*_ *n*_*j*_ is described by the rates of eq. 2. It is important to notice that in this model, contrarily to standard models in population genetics that assume a fixed population size *N*, both the relative abundances of each strain *n*_*i*_*/N* and the total population size *N* = ∑_*i*_ *n*_*i*_ are changing stochastically over time in an interdependent manner. Even at stationarity, the population size *N* is not fixed but oscillates around a typical value, determined by the strains’ parameters.

The presented framework includes all the works based on a competitive Lotka-Volterra like evolutionary dynamics [28– 34] (for *β*(*z*) = 1 and *ω*(*z*) = *z*), and generalization of the Moran model [35] (for *β*(*z*) = 1 and *ω*(*z*) = *α*(1 + *z*)).

## III. STOCHASTIC DYNAMICS

While we want to include stochasticity in our description, taking into account the effect of finite population sizes, we are interested in the limit of large population sizes, which in our context corresponds to *K*_*i*_ *»* 1. In this limit classic models of population genetics (such as the Moran model or Wright-Fisher model) converge to the same effective diffusive limit, which corresponds to Kimura’s equation.

The birth-death process defined in eq. 5 can be approximated by Itô stochastic differential equations as follows

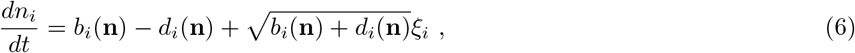

where *ξ*_*i*_ is a Gaussian white noise, ⟨ *ξ*_*i*_(*t*) ⟩ = 0, ⟨ *ξ*_*i*_(*t*)*ξ*_*j*_(*t*′) ⟩ = *δ*_*ij*_*δ*(*t − t* ′) (see Appendix D for the derivation).

### A. Separation of ecological and evolutionary timescales

Eq. 6 describes the stochastic dynamics in the limit of large population sizes (*K*_*i*_ *»* 1). In addition to this limit, we consider the limit of small fitness differences (|*ϕ*_*i*_ *− ϕ*_*j*_|*/ϕ*_*i*_ *«* 1). In this case, as depicted in Figure 3, the dynamics separate in a fast transient followed by slow dynamics (see Appendix E). The initial trajectory drives the system to a slow manifold of solutions, corresponding to the equilibria of the deterministic dynamics in absence of fitness differences. What follows is a dynamic constrained on the slow manifold.

**FIG. 3:**
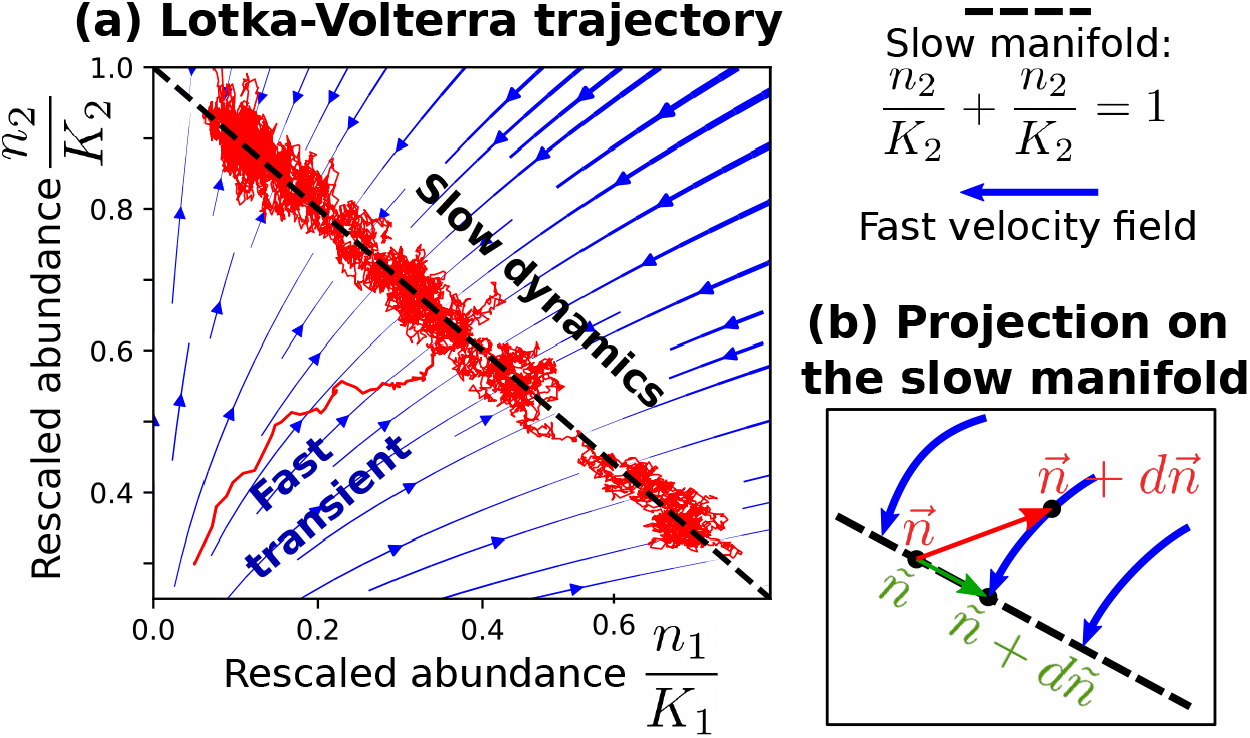
The panel (a) shows an illustrative example of the time-scale separation in a Lotka-Volterra for two species: *β*(*z*) = 1, *ω*(*z*) = *z, K*_1_ = *K*_2_ = 2000, *t*_1_ = *T*_2_ = 1, *ϕ*_1_ = 1, *ϕ*_2_ = 1.002. The variables *n*_*i*_*/K*_*i*_ are the abundances re-scaled by the carrying capacity. The panel (b) sketches the idea underlying the projection of the dynamics onto the slow manifold: infinitesimal steps outside the manifold (in red) are mapped again on it by the fast velocity field. As a result, to correctly take into account these effects, the increment of the effective variable *ñ* (in green) corresponds to the projection of 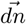 along the velocity field.

It is convenient to describe the dynamics on the slow manifold in terms of the relative abundances *x*_*i*_ = ∑ *n*_*i*_*/* _*j*_ *n*_*j*_. The replicator equation 4 describes the deterministic dynamics of the relative abundances on the slow manifold only under the specific parameter choices discussed in sec. II C. Its derivation does not trivially generalize to the stochastic case and in presence of fully general parameter combinations. The “slow” dynamics is in fact determined by the combination of two forces (see Figure 3). Stochasticity (through genetic drift) pushes the system away from the manifold of solutions. The deterministic part of the dynamics is pushing the system back to the manifold. These two forces do not act orthogonally to the manifold, but the non-linearity of population dynamics and the multiplicative nature of demographic stochasticity result in a non-trivial combination with a net force, which corresponds to an effective frequency-dependent selection that drives the evolution of the system.

In the following we consider two strains, where *x ≡ x*_1_ as the relative abundance of strain 1 and *x*_2_ = 1 *− x*. The effective dynamics on the slow manifold reduces to

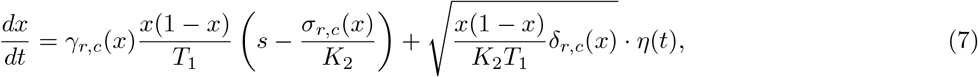

where *η*(*t*) is a white noise term (see Appendix E). In addition to *K*_2_ and *T*_1_, the dynamics of the relative abundance *x* depends on three parameters, *s, r, c*, through the three functions *γ*_*r,c*_(*x*), *σ*_*r,c*_(*x*), *δ*_*r,c*_(*x*) specified below. These three quantities are related to the ratios between the parameters characterizing the population dynamics of the two strains.

In particular, *s* is the selection coefficient *s* = (*ϕ*_1_ *− ϕ*_2_)*/ϕ*_1_ while *r* = *T*_2_*/T*_1_, and *c* = *K*_2_*/K*_1_. The three functions read

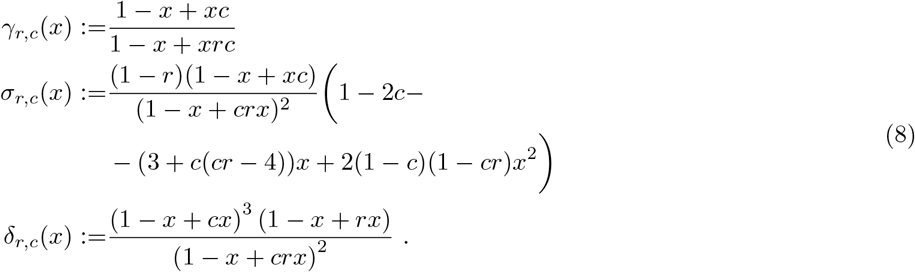

The trajectories of the relative abundances of two strains growing together are therefore determined also by differences in inter-generation times (case *r /*= 1) and in carrying capacities (case *c* 1). If the inter-generation times *T* and the carrying capacities *K* are equal (i.e., if *r* = *c* = 1), eq. 7 reduces to Kimura’s diffusion limit. In fact, *γ*_1,1_(*x*) = *δ*_1,1_(*x*) = 1 and *σ*_1,1_(*x*) = 0.

The sign of the deterministic term of equation 7 depends on the balance between two forces. The first term is the selection coefficient *s*. The second one (equal to *σ*_*r,c*_*/K*_2_) is absent in the classic Kimura’s diffusion limit. It depends non-trivially on the time scales, the carrying capacities, and strain frequencies.

To better understand this dependence, it is instructive to consider the growth of a rare strain with relative abundance Since the strain is rare the initial growth (or decline) will be approximately described by a stochastic exponential growth, which can be obtained by expanding eq. 7 for *x «* 1. In particular, the average relative abundance *x* is determined by

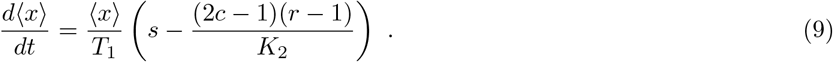

If the carrying capacity of the resident population is very large, *K*_2_ *→ ∞*, the growth rate converges to the deterministic limit *s/T*_1_, depicted in Fig. 2. More precisely, the deterministic limit is correct whenever *sK*_2_ *»* (1 *−* 2*c*)(1 *− r*).

On the other hand, the presence of large but finite population sizes and large enough trait variation drastically change the relative abundance trajectories. A particularly interesting scenario emerges when 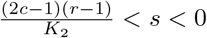. In that case, accordingly to standard results strain 2 would a selective advantage over strain 1 (*s <* 0). However, the presence of the other extra-term counterbalances the positive value of the selection coefficient, making strain 1 able to invade. This implies that, under some choices of *c* and *r*, it can happen that an invader having smaller invasion fitness can grow on average within the “fitter” resident population. To better quantify this statement, in the next section we compare the fixation probabilities of the two strains.

## IV. CONSEQUENCES ON EVOLUTIONARY DYNAMICS

### A. Carrying capacity, inter-generation times, and invasion fitness control the evolutionary success

The evolutionary success of a strain in an environment described by 7 can be quantified by the fixation probability: the likelihood that one individual of a given strain is able to invade a population of a second strain. The ratio between the probability of fixation of one individual of the first strain *p*_1fix_ in a resident population of strain 2 and the fixation of one individual of the second strain *p*_2fix_ in a resident population of strain 1 can be expressed analytically (see Appendix F) and equals to

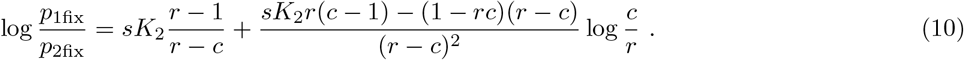

which, in the singular case of *r* = *c* becomes log 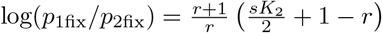.

Despite the generality of the modeling framework 5, this quantity depends on only three parameters: the inter-generation times ratio *r*, the ratio between the carrying capacities *c*, and the product between the selection coefficient and the size of the second population at carrying capacity *sK*_2_.

Figure 4a shows the surface log(*p*_1fix_*/p*_2fix_) = 0, which separates the regions of parameters between a more successful first strain, *p*_1fix_ *> p*_2fix_, and the opposite scenario. Note that, differently from the deterministic approximation of eq. 3, invasion fitness does not unequivocally determine the most likely evolutionary outcome. It exists in fact a region of parameters within the volume below the *s* = 0 plane and above the log(*p*_1fix_*/p*_2fix_) = 0 surface of the Figure 4 for which a the strain with negative selection coefficient is more likely to invade than the one with higher invasion fitness. In absence of differences in the inter-generation time and carrying capacity (i.e., if *r* = *c* = 1), one recovers the Kimura’s diffusion limit and the classical result obtained under a constant total population size *K*_2_ [8]: log(*p*_1fix_*/p*_2fix_) = *K*_2_*s*

**FIG. 4:**
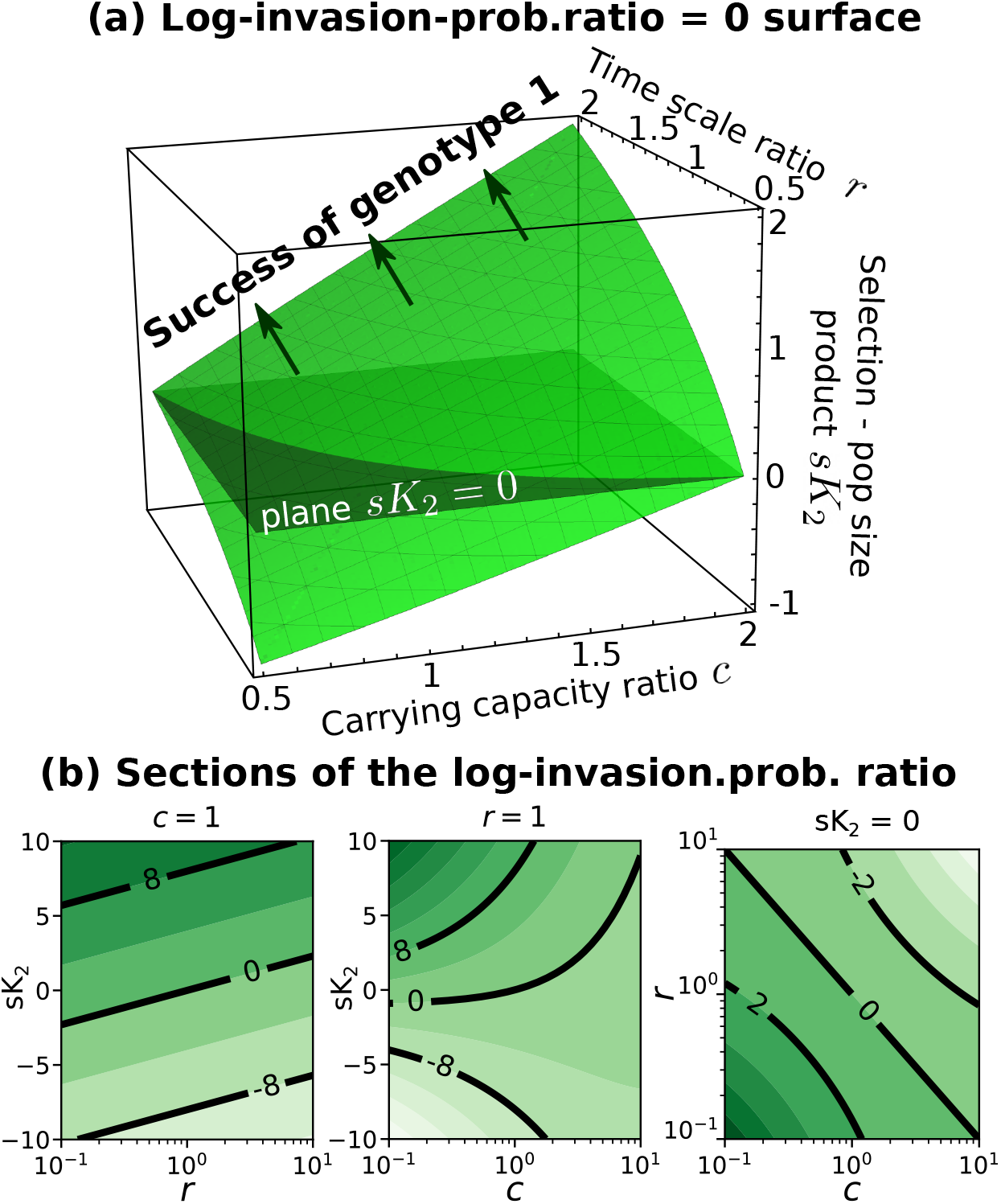
How the fixation probability depends on the three parameters. Panel (a) shows the surface log(*p*_1fix_*/p*_2fix_) = 0, panel (b) different sections of the fixation probability log-ratio. A darker shade of green represents a larger ratio.

It can also be instructive to isolate the dependency on the carrying capacities when intrinsic fitnesses and inter-generation times do not vary (i.e., if *s* = 0 and *r* = 1). In that case, *p*_1fix_*/p*_2fix_ = 1*/c* = *K*_1_*/K*_2_. This implies that that larger carrying capacities, corresponding to lower demographic fluctuations and genetic drift, are favoured. Similarly, if the selection coefficient equals 0 and the carrying capacity are equal between the strains (*c* = 1), the ratio of the fixation probabilities reduces to *p*_1fix_*/p*_2fix_ = 1*/r* = *T*_1_*/T*_2_. Also in this case, longer inter-generation times, which correspond to lower per-capita birth and death rates at carrying capacities, and therefore lower stochasticity, are favored.

### B. The evolutionary outcome is independent of modeling details

One key result of our derivation is that the evolutionary trajectories, described in the diffusion limit by eq. 7, do not depend on the specific modeling choices. More precisely, eq. 7 does not depend on the shape of the functions *β*(*z*) and *ω*(*z*).

Figure 5 shows numerical simulations of the fixation probability in alternative models, differing for the mathematical dependency of birth and death rates on strain abundances. The figure shows that once these different models are mapped onto our framework 5, all the invasion probabilities collapse onto the same curves as predicted by equation 10. The results of figure 4 are therefore general and do not depend on the specific model considered. For instance, for *c* = 1 and *s* = 0 one recovers the fixation probability of [35] and [33]. Details about how the simulations are performed are shown in Appendix G. The definitions of the simulated models and how to map them in our framework are in Appendix A.

**FIG. 5:**
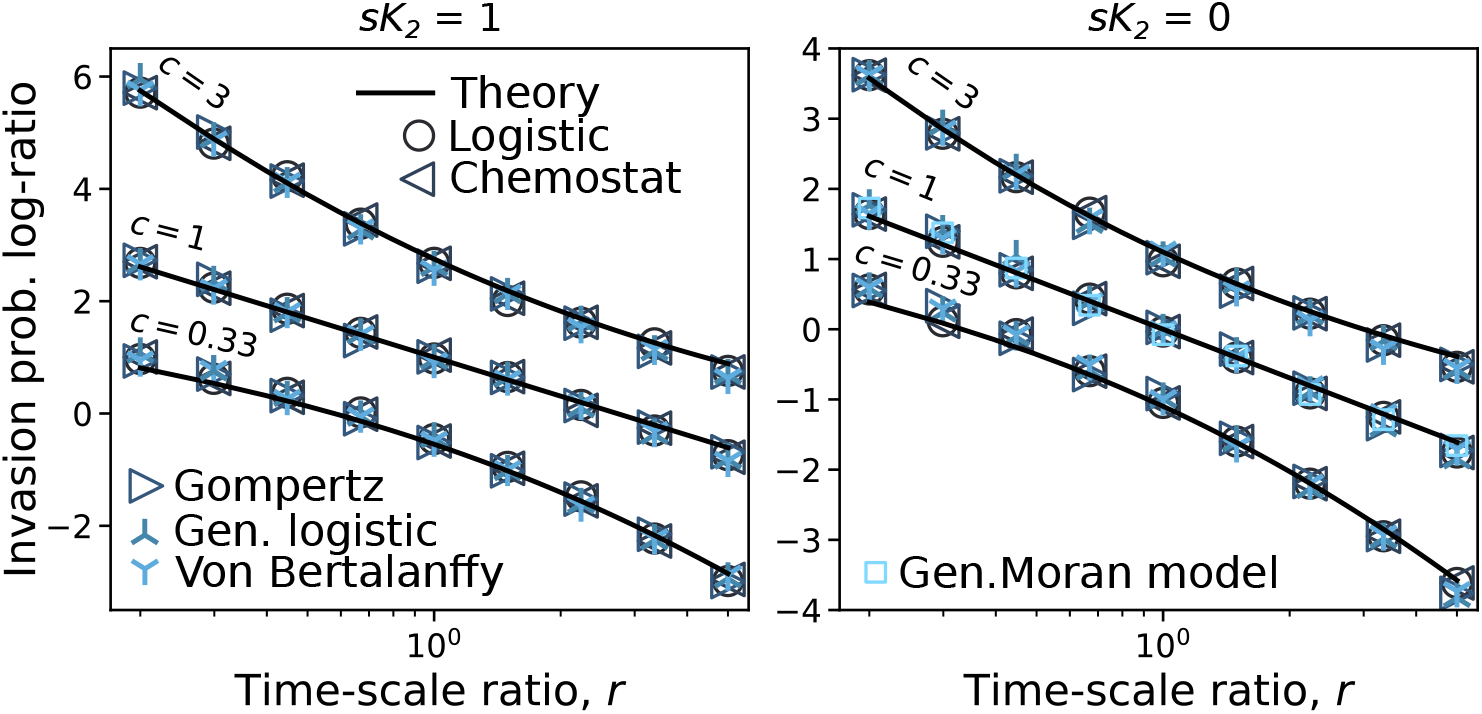
Collapse of competitive models on the same fixation probability, given by equation 10, black lines. The simulations consider the competitive Lotka-Volterra, a chemostat model [42], the Gompertz growth dynamics [43, 44], a generalized Lotka-Volterra [45] with exponent *ν* = 1*/*2, the Von Bertalanffy model [46] with *α* = 2*/*3 and the Generalized Moran model [35] which is defined only for *sK*_2_ = 0 and *c* = 1. For more details about the models see Appendix A *K*_2_*s*.

### C. The asymmetry in total mutation rates induces an additional dependency on population size and inter-generation timescale

The expected number of mutations per generation depends on the population size. Since in our case the total population size can vary depending on which strains are present and in what abundance, also the total mutation rate depends on which strains are present and in which abundance. This dependency of the total mutation rate on population size induces an additional effect of the varying total population on the most likely evolutionary trajectory. Let us consider two strains and a symmetric mutation probability *U «* 1. If we start with a clonal population of strain 2 at carrying capacity *K*_2_, the birth and death rates of 5 are balanced by definition of steady state. The birth rate reads therefore *b*_2_ = *ρ*_2_*β*(*ω*^*−*1^(*ϕ*))*K*_2_. By multiplying this rate by the mutation probability *U* one obtains the mutation rate, which, in this simple case, corresponds to the rate at which an individual of strain 1 appears. Therefore, one can compute the rate at which strain 1 substitutes the resident population composed of strain 2, which equals *m*_1fix_ = *Ub*_2_*p*_1fix_. The ratio of this quantity for the two strains reads then

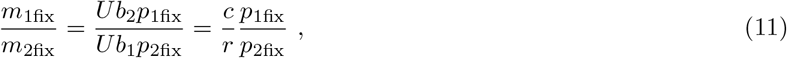

where the ratio *p*_1fix_*/p*_2fix_ is given by eq. 10. Including explicitely the effect of generation time and population size on mutation introduces an additional dependency on *c* and *r*.

In the periodic selection regimes, i.e., when the time between two successful mutations is much longer than the time to fixation of a successful mutation, this ratio is directly related to the relative probability of finding the population dominated by strain 1 vs strain 2.

Figure 6 shows the analytical behavior of this expression. Also in this case there is a large range of parameters for which the population evolves to lower values of invasion fitness.

**FIG. 6:**
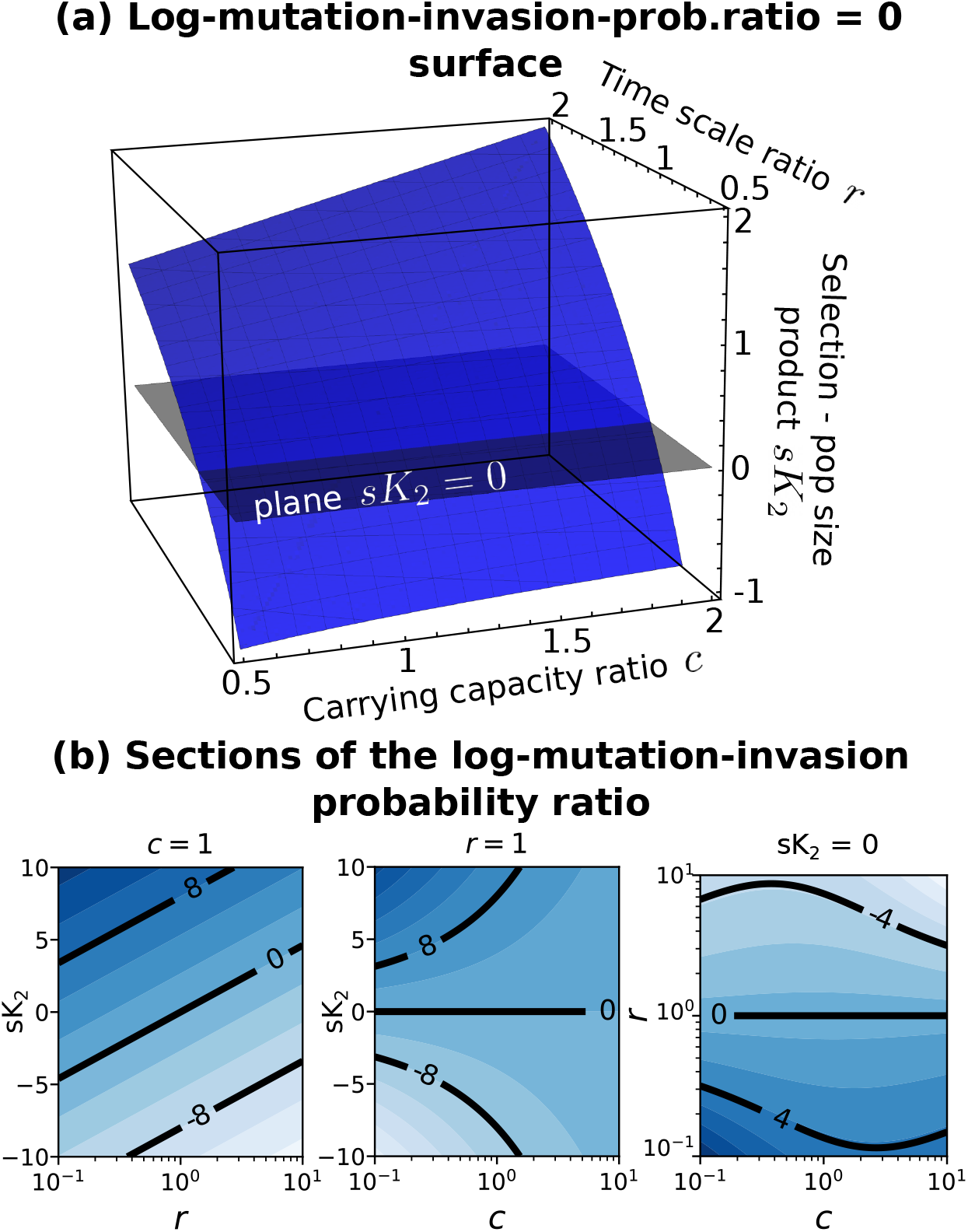
Behavior of the invasion probability through mutations. The blue surface is log(*m*_1fix_*/m*_1fix_) = 0 defined in eq. 11

### D. Evolution in the chemostat model with metabolic trade-off can revert the direction of evolution

As an example for the application of the previous results, we consider a population growing in a chemostat on externally provided resources [36]. We consider a birth-death process where the birth-rate depends on the expected availability of resources, while the death rate is a constant factor. In the deterministic limit the equation reduces to

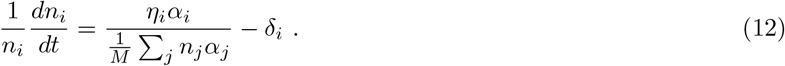

The parameter *α*_*i*_ represents the resource intake rate of strain *i, δ*_*i*_ is a death/dilution rate, while *Mη*_*i*_ is the efficiency of resource-to-biomass conversion (equivalent to an inverse yield), and *M »* 1 a parameter that sets the scale of the total population size. One can then map this model into our general framework, for the choice *β*(*z*) = 1*/z* and *ω*(*z*) = *z*, as described in Appendix A

A particularly interesting scenario appears when *α, δ*, and *η* are not independent, being subject to a tradeoff. For instance, yield (or, equivalently, efficiency) decreases during selection in experimental evolution [47–49], giving rise to a trade-off between growth rate and yield.

Specifically, we focus on mutations of the resource intake rate *α* and consider the case *δ*_*i*_*/η*_*i*_ = *α*_*i*_ + *α*_0_, where *α*_0_ *«* 1 is a small parameter (in principle of the order of 1*/M*). This choice implies that the population size at carrying capacity in a clonal population, i.e. *K*_*i*_ = *Mη*_*i*_*/δ*_*i*_ = *M/*(*α*_*i*_ + *α*_0_), decreases with the intake rate *α*, giving rise to the growth-rate and yield trade-off.

In the deterministic limit, it is easy to see that strains with higher intake rates (i.e. with higher invasion fitness *ϕ*_*i*_ = *α*_*i*_*η*_*i*_*/δ*_*i*_ = *α*_*i*_*/*(*α*_*i*_ + *α*_0_)) always invade populations with lower values of *α*. By using using equations 10 and 11, we show that the presence of a large, but finite, strain-dependent total population size can invert this behavior. Moreover, the outcome of the evolutionary process depends on whether the dependency on *α* is included in the death rate or in the efficiency. We consider here the former case, while we discuss the latter, which nevertheless leads to similar results (see Appendix H).

If the intake conversion from the resource to the biomass is constant, *η*_0_, the death rate per-capita can be expressed as *δ*_*i*_ = *η*_0_(*α*_*i*_ + *α*_0_). The computation of the time-scale ratio leads to *r* = *c*. In such a case, there is no difference between external invasions, eq. 10, or invasions through mutations, eq. 11. In particular, using eq. F9 (the special case *r* = *c* of eq. 10) one obtains

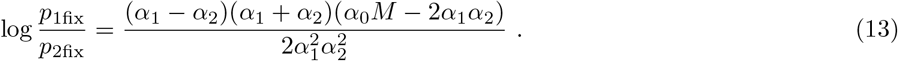

Strain 1 is advantaged (*p*_1fix_*/p*_2fix_ *>* 1) only if *α*_0_*M −* 2*α*_1_*α*_2_ *>* 0. If, without loss of generality, we assume *α*_1_ *> α*_2_, we obtain that strain 1 is advantaged over strain 2 in the case *α*_2_ *< α*_1_ *< Mα*_0_*/*(2*α*_2_). Therefore if a strain *i* has a value of intake rate *α*_*i*_ such that *α*_*i*_ = *Mα*_0_*/*(2*α*_*i*_), no other strain can be advantaged over it. This indicates the existence of an optimal value of *α*, equal to 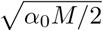, to which the evolutionary trajectories converge.

Figure 7 shows a simulation of this scenario, where *α* can mutate taking discrete, uniformly spaced, values *{α*_*i*_*}*. The evolutionary trajectory, starting both above and below the predicted optimal value converges to it in the long term. Figure 7b shows that the expected value of the intake rate *α* (and therefore the invasion fitness) decreases over time if the population has initial values of *α* above the predicted value.

**FIG. 7:**
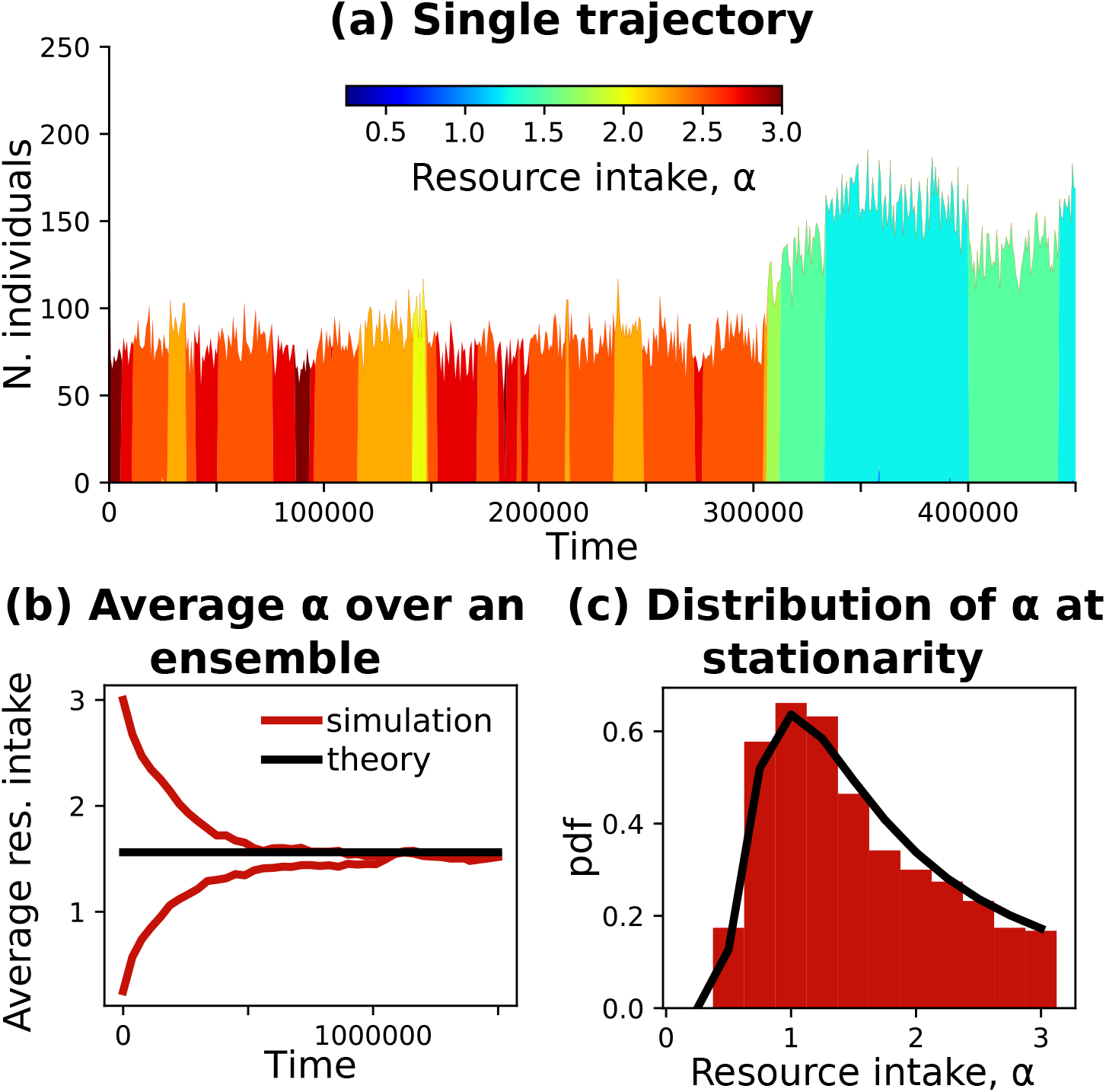
Evolutionary dynamics of the chemostat with a constant intake rate *η*_0_ = 1. The resource intake *α* can mutate to a neighbour *α* + Δ*α* or *α −* Δ*α* with rate *U* = 2 *·* 10^*−*5^. It is constrained to be in [0, 3]. The other parameters are: Δ*α* = 0.25, *α*_0_ = 0.01, *M* = 200. Panel (a) shows the number of individuals colored by their value of *α* of a Gillespie simulation. There is no clonal interference since, typically, there is no co-existence of sub-populations with different *α*. Panel (b) is the average resource intake over 300 realizations of two ensembles starting from different initial conditions. Panel (c) is the distribution of *α* at stationarity of the 300 realizations.

The presence of stochasticity implies that the optimal value is reached on average, with a fluctuating value of *α* around the optimal value. Under the assumption of periodic selection, one can predict the distribution *p*(*α*) at equilibrium by assuming that evolution follows a jump process which satisfies the detailed balance: *p*(*α*_*i*_)*p*(*α*_*i*_ *→ α*_*i*+1_) = *p*(*α*_*i*+1_)*p*(*α*_*i*+1_ *→ α*_*i*_) (details in Appendix H). The prediction for the average and the full distribution is in agreement with the simulation in panels (b) and (c) of Figure 7.

## V. DISCUSSION

Here we have studied the evolutionary trajectories of populations in a broad class of models characterizing population self-limitation. We introduced a population dynamics framework that encompasses several known models (Logistic, Gompertz, Chemostat, and others). In presence of large, but finite, population sizes, we have shown that the evolutionary trajectories depend only on three quantities (timescale, invasion fitness, and carrying capacity), irrespective of the models’ specific form.

We obtained these results by assuming a time-scale separation between the total population size and the strain (or alleles) frequencies [28, 31, 33, 35]. This assumption allows us to write down an effective equation that describes the time evolution of strain frequencies, which depends only on the values of the three evolutionary relevant quantities. This effective description reduces to Kimura’s diffusion limit when strains differ only in their invasion fitness or in the limit of strong selection. In that case, variation in timescales and/or carrying capacities become effectively irrelevant. One especially interesting aspect of the effective equation we obtained is the role played by the (finite) population size. In Kimura’s diffusion limit, population size influences only the strength of genetic drift. While in that context drift masks the effect of selection (e.g., a deleterious mutation has a non-zero fixation probability), it never alters its direction (i.e., a beneficial mutation has always a larger fixation probability than a deleterious one). In our setting, the existence of finite population size might alter the course of evolution: mutants with lower invasion fitness might be more likely to invade than strains with a higher invasion fitness.

We explicitly show this effect in the chemostat model in the presence of a metabolic tradeoff. In the deterministic limit, larger resource intakes correspond to higher fitness, and therefore evolution drives the population to higher and higher resource intakes. In presence of a large, yet finite, population size, the naive expectation obtained in the deterministic case is not realized. The evolutionary trajectory converges in fact to an optimal value of the intake rate, which we analytically predict. If a clonal population has an initial intake rate larger than the optimal one, evolution will drive the population to decrease the intake rate, and therefore invasion fitness.

Invasion fitness decreases over time because of the presence of a finite population size. This might be reminiscent of Muller’s ratchet [50–52], but it has a radically different origin. In the case of Muller’s ratchet, the decrease in fitness is determined by the fact that mutations only give rise to deleterious mutations, which can get fixed because of the finite population size. The presence of a small fraction of beneficial mutations is enough to balance the effect of deleterious mutations and lead to an overall increase in fitness [53].

In our case, the mechanism in play is radically different. The presence of variation in the timescale and/or in the carrying capacity creates an effective force driving the population to higher values of the carrying capacity and lower values of the inter-generation times. The parameter identified as invasion fitness in the deterministic dynamics does not predict the (likely) outcome of the evolutionary dynamics. Lower carrying capacities correspond to a higher level of genetic drift (as shown in eq. S12 of the Supplementary Materials). The intuition is that the evolutionary trajectory drives the population toward lower values of genetic drift. This effect is alike to thermophoresis [54] or to what is observed in Brownian motion in presence of a position-dependent diffusion: an effective force drives the trajectories to lower values of noise [55]. In our case, the effective force can counterbalance the sign of the selection coefficient.

This result highlights the relevance of genetic drift in shaping evolutionary trajectories. In our case, drift not only affects the speed of evolution [56], but also its direction. One remarkable aspect is that this effect turns out to be model-independent and determined only by differences in the timescale and carrying capacity.

The three quantities (invasion fitness, carrying capacity, and timescale) that effectively determine the trajectory of an evolving population are not independent traits. Variation of a given trait can influence one, two, or all three of them. For instance, in the example of the chemostat model that we discussed, only one trait, the resource intake, is under selection pressure. But since it influences all three quantities, the result of evolution is far from trivial.

Our results could be reminiscent of r/K selection [57, 58], but the connection is not obvious. In saturated environments, i.e. when populations are close to carrying capacities, K-selection dominates, favoring strains with high-competitive abilities (which in our setting correspond to higher values of *ϕ*). In un-saturated environments, r-selection favors strains with higher fecundity (in our setting, lower values of *T*). We considered the case of a saturated environment, studying evolution when total population abundance was close to carrying capacity. The effect of demographic stochasticity, as opposed to environmental fluctuations at the core of r/K selection, was responsible for non-trivial evolutionary effects (e.g., the existence of an optimal intake rate in the chemostat model) which disappear in the limit of large population sizes.

Our results go in the direction of building a model-independent eco-evolutionary theory. The extent to which the specific details of ecological interactions influence the evolutionary outcome is a major limitation to the development of a comprehensive general understanding of eco-evolutionary trajectories. Our results present a first step in this direction, as they describe the evolution of a population limited by a single factor. The next step will be to generalize our framework by considering multiple limiting factors, giving rise to the coexistence of different populations [59]. As a further generalization, one can consider different interaction types, going beyond the competition.

One important hypothesis of our analysis was to consider the case of strain independent functional forms of *ω*(·) and *β*(·). This is a key assumption to obtain model-independent evolutionary trajectories. One could consider the more general scenario where the functional forms *β*_*i*_(·) and *ω*_*i*_(·) explicitly depended on strain *i* in a way that could not be recast to the one eq. 3. While this case could be interesting for future work from a theoretical standpoint, it is important to highlight that all the commonly used models could be written in terms of eq. 3 and did not require introducing strain-dependant functional forms. Another limitation of our framework, again related to the choice of *ω*(·) and *β*(·), lies in the fact that we restrict our analysis to a class of models that admit a Lyapunov function, through the hypothesis of monotonic *ω*(·). This framework does not allow to include mechanisms allowing multi-stability (e.g. Allee effect).

In this work, we have only considered haploid populations. The extension to diploid (or complex) traits is numerically easy to implement as our setting can be generalized fairly simply. It is much more complicated to obtain closed equations for the diffusive limit under a separation of time scales. Preliminary analysis of numerical examples, shows however that, also in the diploid case, evolutionary trajectories are model-independent.

To conclude, our work shows that several aspects of the population dynamics are details that do not influence the trajectories of an evolving population. Only three demographic parameters effectively matter. Their variation is subject to selection in a non-trivial, drift mediated, way, which might lead to a decrease in what is usually identified as fitness over time.

## Appendix A: Definition of the ecological models

### 1. Logistic growth and competitive Lotka-Volterra

Logistic growth is recovered for *ω*(*z*) = *z* and *β*(*z*) = 1. In this case, the per-capita growth rate of a clonal population, obtain from eq. 1 reads

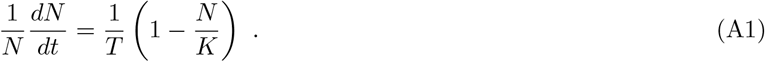

In the case of multiple strains, eq. 3 we obtain

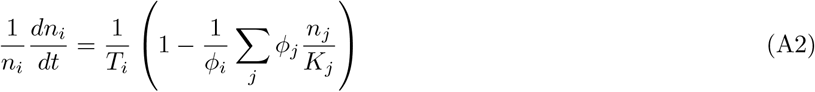

### 2. Chemostat

To derive the chemostat dynamics, let us first consider a clonal population. Let us assume that the per-capita birth rate depends on the concentration *R* of some resource, *b*(*N*)*/N* = *αηH*(*R*), where *H*(·) is some monotonic function of the concentration, an intake-rate *α*, and a conversion factor between resource intake and biomass *η*. We assume that the per-capita death rate is independent of abundance *d*(*N*)*/N* = *δ*. The resource concentration changes accordingly to a classic chemostat equation

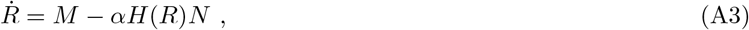

where the first term represents a constant influx of resources and the second term is consumption. We assume that the resource concentration *R* quickly reaches a stationary value, which leads to *H*(*R*^*∗*^) = *M/*(*Nα*) at the stationary value. We obtain therefore that *b*(*N*) = *ηM*, from which we get

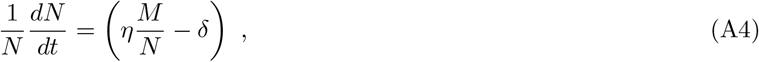

which corresponds to the case *β*(*z*) = 1*/z* and *ω*(*z*) = *z*.

This can be easily extented to multiple strains, following [42] in the case of a single resource, by assuming that *α*_*i*_, *η*_*i*_ and *δ*_*i*_ have strain-dependent values, while resource concentration changes as

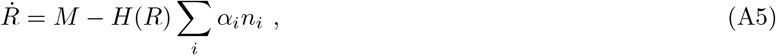

from which, after a time-scale separation, we obtain eq 12.

This model maps in our general framework using *T*_*i*_ = 1*/*(*α*_*i*_*η*_*i*_), *ϕ*_*i*_ = *α*_*i*_*η*_*i*_*/δ*_*i*_, and *K*_*i*_ = *Mη*_*i*_*/δ*_*i*_.

### 3. Gompertz growth

Gompertz growth [44] of a clonal population is defined as

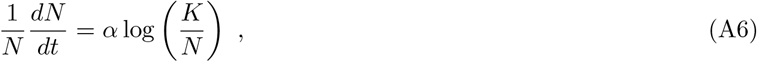

which corresponds to *β*(*z*) = 1 and *ω*(*z*) = log(*z*).

We generalize Gompertz growth to the case of multiple strains as

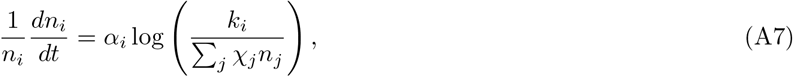

which recovers the classical Gompertz growth dynamics in the case of a single strain and *χ*_1_ = 1. The mapping with the general framework is obtained uwing: *T*_*i*_ = 1*/*(*α*_*i*_ log(*k*_*i*_)), *ϕ*_*i*_ = log(*k*_*i*_), and *K*_*i*_ = *k*_*i*_*/χ*_*i*_.

### 4. Generalized Lotka-Volterra

The generalized logistic dynamics [45] in a clonal population is defined as

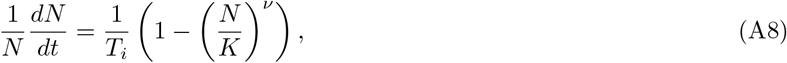

where *ν* is a positive exponent. The generalize logistic dynamics correspond to the choice *β*(*z*) = 1 and *ω*(*z*) = *z*^*ν*^

The case of multiple strains can be written as follows:

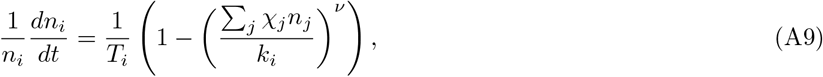

whose mapping reads: 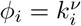, and *K*_*i*_ = *k*_*i*_*/χ*_*i*_. The simulations of figure 5 are obtained for *ν* = 1*/*2.

### 5. Von Bertalanffy model

The Von Bertalanffy growth [46] is defined by the equation

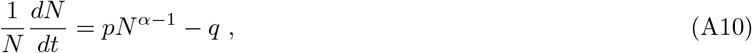

where *α ∈* [0, 1). Von Bertalanffy growth corresponds to *β*(*z*) = *z*^*α−*1^ and *ω*(*z*) = *z*^1*−α*^.

The natural extension to competing strains reads:

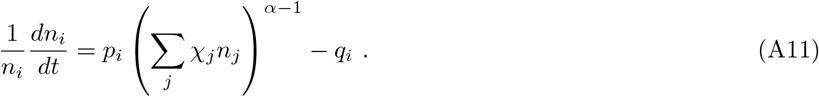

It is included in the general framework by choosing *T*_*i*_ = 1*/p*_*i*_, *ϕ*_*i*_ = *p*_*i*_*/q*_*i*_, and *K*_*i*_ = (*q*_*i*_*/p*_*i*_)^1*/*(*α−*1)^*/χ*_*i*_. The simulations of figure 5 are obtained for *α* = 2*/*3.

### 6. Generalized Moran model

The Generalized Moran model [35], is defined by the birth and death rates

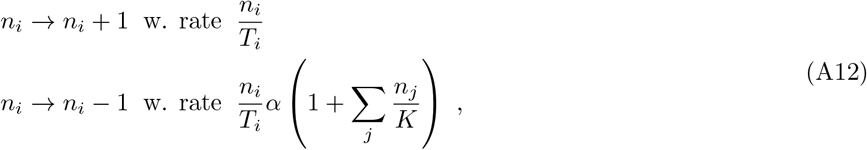

which corresponds to the case *β*(*z*) = 1 and *ω*(*z*) = *α*(1 + *z*), where *α* is the same across strains. Note that, in this case, *K*_*i*_ = *K* and *ϕ*_*i*_ = 1 for all strains.

This corresponds in fact to the “quasi-neutral” case, where *α*, defined as the ratio of death and birth rate is constant for each genotype.

## Appendix B: Deterministic dynamics of multiple strains

Equation 3 describes the deterministic limit of the dynamics in the case of multiple strains. A Lyapunov function for the dynamical system of eq.3 reads

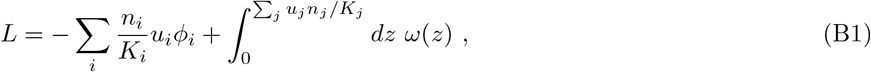

where *u*_*j*_ = *ω*^*−*1^(*ϕ*_*j*_). One can in fact show that

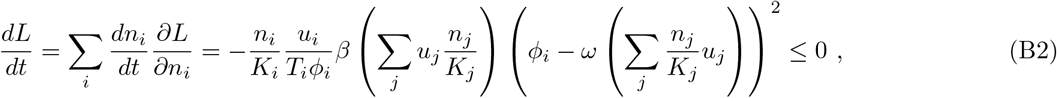

under the assumption that *β*(·*·*) is positive. The second derivatives of *L* read

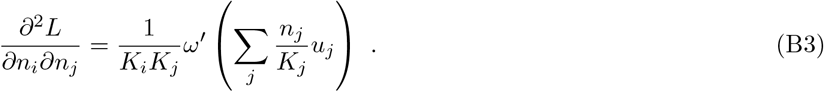

The Hessian is therefore positive definite if and only if the function *ω*′ (·*·*) is monotonically increasing, i.e. if *ω* (·) is a positive function. Under this assumption, the function *L* is a convex function. The fact that the function *ω*(·) is monotonically increasing also implies that the function defined in eq. B1 is bounded from below.

The existence of a convex Lyapunov function implies that, in the non-degenerate case (i.e., where multiple populations are characterized by the same value of *ϕ*) there is a unique, globally stable, fixed point of eq. 3. By minimizing eq. B1, one obtains that the globally stable fixed point is

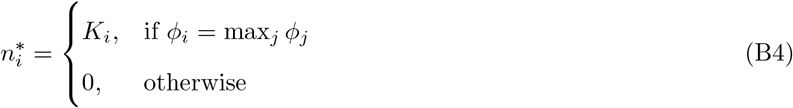

Unless multiple strains are characterized by the same values of *ϕ*_*i*_, eq. 3 does not, therefore, allow the coexistence of multiple strains. For large times, the community is dominated by the strain with the largest value of *ϕ*_*i*_.

## Appendix C: Interpretation of parameters

The parameter *K*_*i*_ represent the carrying capacity of a strain in isolation. In absence of others, in fact, the birth and death rate of a strain (eq. 5) are equal to each other when *n*_*i*_ = *K*_*i*_. In particular, they reads

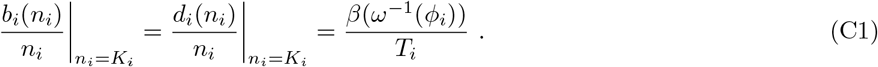

Up to the factor *β*(*ω*^*−*1^(*ϕ*_*i*_)), the parameter *T*_*i*_ represents therefore the expected generation time, i.e. the average interval between the birth of an individual and the birth of its offspring, when the population is at carrying capacity.

For instance, in the case of the logistic growth (when *β*(*z*) = 1), *T*_*i*_ exactly corresponds to the generation time at carrying capacity.

Moreover, when a clonal population is rare *n*_*i*_ *« K*_*i*_, the average growth of strain *i* reads

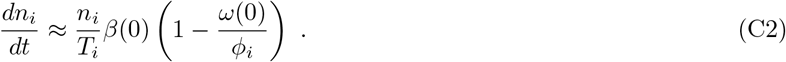

The population is therefore expected to grow exponentially, with growth rate proportional to 1*/T*_*i*_. The proportionality constant is model dependent, and in particular depends on the values *ω*(0) and *β*(0). For instance, in the Logistic case, since *ω*(0) = 0 and *β*(0) = 1, the growth rate is exactly equal to 1*/T*_*i*_. Note that for some of the models, *β*(*z*) and/or *ω*(*z*) diverge in the limit *z →* 0. In those cases, the growth at small populations abundances is not exponential.

As shown in Appendix B, the values of *ϕ* unequivocally determine which strain survives in the long term, and can therefore be naturally interpreted as fitness. Such an interpretation becomes more clear by considering a rare mutant/invader spreading in the population in presence of a large resident population at carrying capacity. Let the strain 1 be a rare invader (i.e. with initial population *n*_1_ *≈* 1 *« K*_1_) and 2 the type of the resident population. Up to small terms (of the order 1*/K*_1_), the equation 3 for the invader reads:

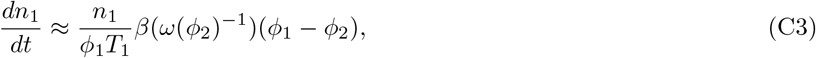

where the growth rate is proportional to (*ϕ*_1_ *− ϕ*_2_)*/*(*ϕ*_1_*T*_1_), as also depicted in Fig. 2.

## Appendix D: Diffusive limit of the birth-death process

Here we are interested in the limit *K*_*i*_ *→ ∞* of the birth-death process 5. In such a limit, the stochastic process can be approximated, via a diffusion limit, with a stochastic differential equation, which treats the abundances *n*_*i*_ as continuous variables. The diffusion limit can be formally obtained by truncating the Kramers-Moyal expansion to the leading terms. Here below we provide an intuitive derivation of the diffusion limit in the case of one strain. The generalization to an arbitrary number of strains does not present any further conceptual difficulty.

Let us consider a very small interval of time Δ*t «* 1*/*(*b*(*n*) + *d*(*n*)) such that it is very unlikely that more than one birth and/or death events occur in Δ*t*. This allows us to write down the probability of having *n* individuals at time *t* + Δ*t* as:

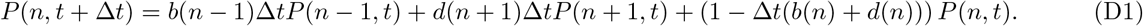

Let us now consider the re-scaled variable *y* = *n/K*, where *K* is the carrying capacity of the strain. The functions that depended on *n ±* 1, now depend on *y ±* 1*/K*. Since *K »* 1 we can consider *y* as a continuous variable and 1*/K* as a small increment, which allows us to expand those functions in the Taylor series. By expanding up to the second term, we obtain

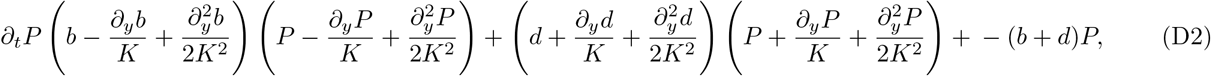

where we omitted the arguments (*y, t*) from all the functions for the sake of a simpler notation, and neglected the terms of order 1*/K*^3^ and higher. By regrouping the terms at different orders of 1*/K*, one obtains

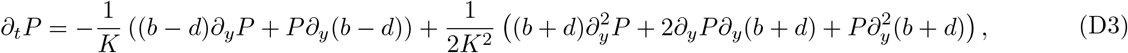

which can be simplified to the following Fokker-Plank equation:

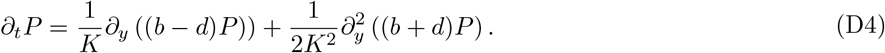

From this expression one can obtain the deterministic coefficient, (*b − d*)*/K*, and the diffusion coefficient, (*b* + *d*)*/*(2*K*^2^), and therefore the associated stochastic differential equation. In the case of *A* strains, the resulting Fokker-Plank reads

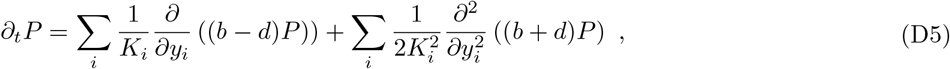

which is equivalent to the following system of Itô SDEs:

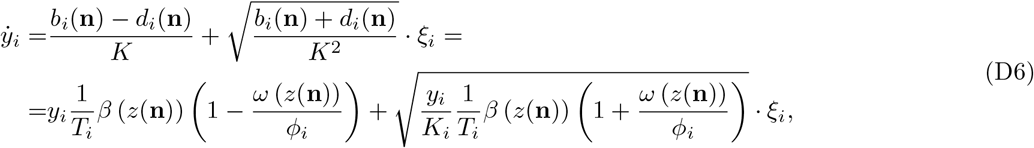

where *ξ*_*i*_ is an uncorrelated normal random variable, ⟨ *ξ*_*i*_(*t*) = 0, *ξ*_*i*_(*t*)*ξ*_*j*_(*t* ′) ⟩ = *δ*_*ij*_*δ*(*t − t*′), and we defined *z*(**n**) =∑ _*j*_ *u*_*j*_*n*_*j*_*/K*_*j*_. From this expression one can easily obtain the equation 6 by simply by change of variable *n*_*i*_ = *y*_*i*_*K*_*i*_. In the limit *K → ∞*, the fluctuations become negligible and one obtains the deterministic limit of equation 3.

## Appendix E: Separation of Ecological and Evolutionary timescales

Let us consider the diffusive dynamics derived above 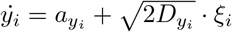, whose coefficients read:

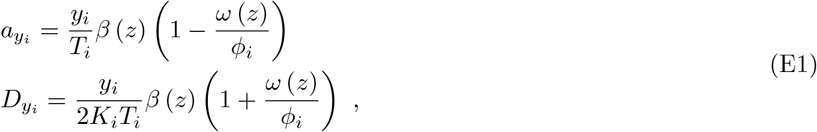

where we have omitted the arguments **n** in *z*(**n**).

When fitness differences are small (i.e., |*ϕ*_*i*_ *− ϕ*_*j*_|*/ϕ*_*j*_ *«* 1) the stochastic dynamics separates in two temporal phases, depicted in Fig. 3. The first phase corresponds to a fast transient: the total population size *N* =∑ _*i*_ *n*_*i*_ changes to reach a value close to the carrying capacity. Fitness differences and stochasticity are negligible in this first transient but determine the dynamics of the second phase.

We introduce the selection coefficient *s*_*i*_ = (*ϕ − ϕ*_*i*_)*/ϕ* of strain *i* relative to strain 1, where *ϕ* = *ϕ*_1_. As we are interested in the limit *s*_*i*_ *«* 1 and *K*_*i*_ *»* 1, we can expand eq. E1. The expansion up to the first first order in *s*_*i*_ and 1*/K*_*i*_ reads,

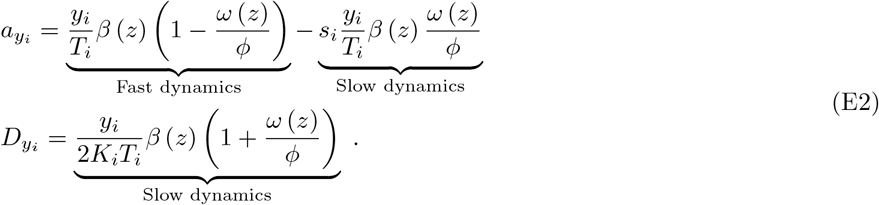

This expansion makes apparent the distinction between fast and slow dynamics. The latter terms are proportional to *s*_*i*_ or 1*/K*_*i*_, which are both small, while the former is not proportional to those factors. The fast dynamics (see Figure 3(a) for an example with Lotka-Volterra dynamics) is determined by the first deterministic term in eq E2, independent of *K*_*i*_ and *s*_*i*_. This fast dynamics drives the system to a manifold of solutions where *ω*(*z*) = *ϕ*, which implies ∑_*i*_ *n*_*i*_*/K*_*i*_ = ∑_*i*_ *y*_*i*_ = 1.

The slow dynamics that follows is determined by the deterministic term linear in *s*_*i*_ and the stochastic term 1*/K*_*i*_.The time-scale separation allows us to consider the effective evolution along the slow manifold. Such effective dynamics is non-trivial (see Figure 3). The small stochastic terms of the original dynamics are no longer negligible and kick the variable outside the manifold. As soon as the abundances move away from the manifold, the fast dynamics become relevant again, and it pushes back — to a non-trivial position — the trajectory onto the manifold.

To correctly take into account this tension between the slow and fast terms, the increment of the effective variable constrained to move along the manifold, can be obtained by projecting the original increment *dy* onto the manifold following the flow field of the fast dynamics. To perform this calculation we take advantage of the fact that there exists a conserved quantity of the fast dynamics. This simplifies a lot the calculation of [60]. The derivation is similar to the computation proposed in [33].

### 1. Fast transient and conserved quantity

The deterministic equation describing the fast transient dynamics reads

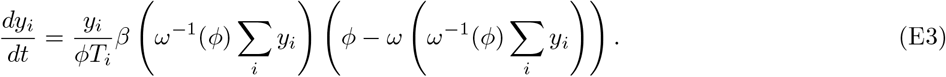

This equation has a manifold of marginally stable fixed points, identified by the simplex ∑_*i*_ *y*_*i*_ = 1. Any point in this simplex is a fixed point of eq. E3. The dynamics drive the system to one of these solutions, which will differ depending on the initial condition.

The existence of a manifold of fixed points is a consequence of a symmetry in the dynamics of eq. E3 which corresponds to the existence of a conserved quantity, which reads

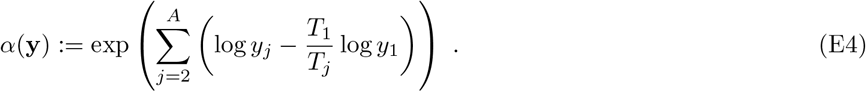

From which one obtains

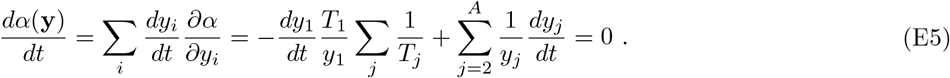

In the simple case of two strains, the conserved quantity reads:

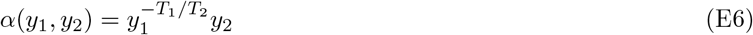

### 2. Effective slow dynamics

The value of the function E4 does not change along the flow field of the fast dynamics and it allows us to correctly calculate the projection onto the slow manifold. In the following, we consider the case of two strains.

Due to the presence of the terms proportional to *s*_*i*_ and 1*/K*_*i*_ in eq. E2 the value of *α*(**y**) is not conserved during the slow dynamics. The time evolution of *α*(**y**) can be obtained from eq. E2 using Itô calculus

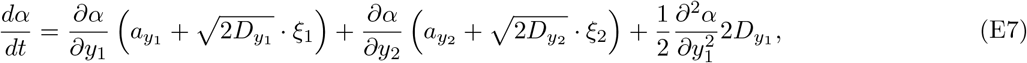

where *a* and *D* are the deterministic and diffusion terms appearing in eq. E2: 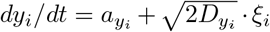. The term with a second-order derivative of *y*_2_ is equal to zero because *α* is linear in *y*_2_. The term with mixed *y*_1_ and *y*_2_ derivatives equals zero because the two noises are uncorrelated. By writing explicitly all the terms, one obtains:

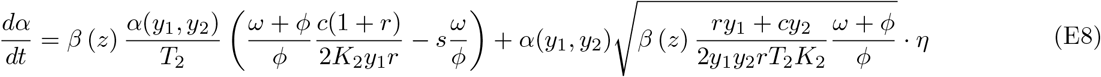

where *η* is a white Gaussian noise, and *r* = *T*_2_*/T*_1_, *c* = *K*_2_*/K*_1_, *s* = *s*_2_ *«* 1.

The key point of the derivation that follows is that the change of *α* outside the manifold is the same on the manifold for a variable projected through the flow field. Let us parametrize the position on the manifold as 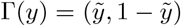, where the variable 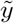 corresponds to *y*_1_ when constrained to be on the slow manifold. The change in *α* for the full dynamics is equivalent to the one obtained by constraining the dynamics on the slow manifold

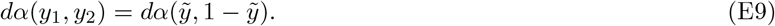

This implies that the dynamics of *α*, evaluated for the variables on the manifold, follow equation E8

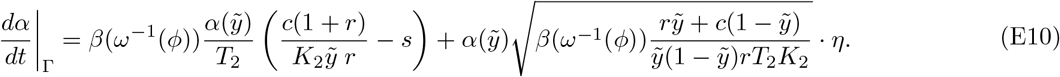

In the following we will remove the term *β*(*ω*^*−*1^(*ϕ*)) as it can always be reabsorbed in the definition of *β*(·).

One can then derive the expression for the re-scaled abundance on the slow manifold 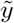 using the Itô formula from eq. E10:

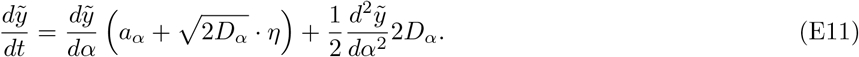

To perform the calculation, one needs to compute the first and second derivatives by using the derivatives of the inverse function *α*(*y*):

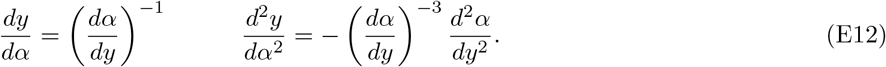

Putting everything together, the final expression for the slow variable is the following:

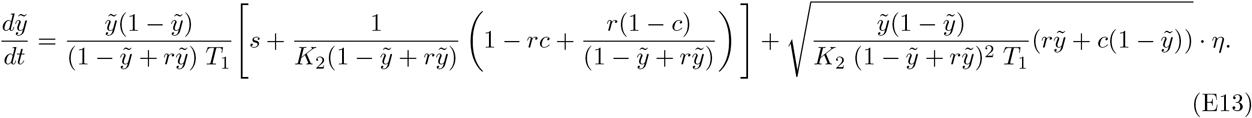

One can express the dynamics in terms of the frequency *x* = *n*_1_*/*(*n*_1_ + *n*_2_), which is related with the re-scaled abundance 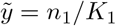 as 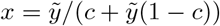. By using again Itô calculus, one finally obtains

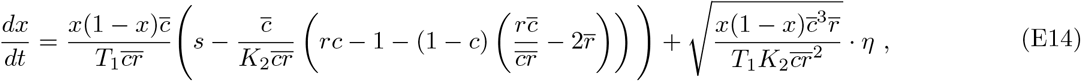

where we used the notation 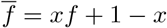 This equation is equivalent to eq. 7.

## Appendix F: Invasion probability

The question that we address now is whether a mutant strain that appears in a population composed of only one resident strain can invade and overcome the original population. To this end, we need the fixation probability 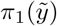, i.e. the probability that, given the initial re-scaled abundance 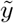, the species will reach 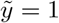 (notice that the same computation can be done starting from the frequencies (E14) leading to the same result).

The fixation probability can be obtained by solving the differential equation 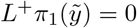, where *L*^+^ is the backward operator of the stochastic process, and the boundary conditions are *π*_1_(0) = 0 and *π*_1_(1) = 1. This leads to the following formula:

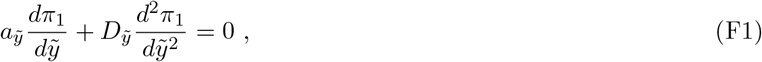

The solution of this equation reads:

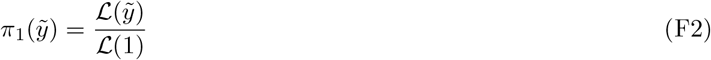

where

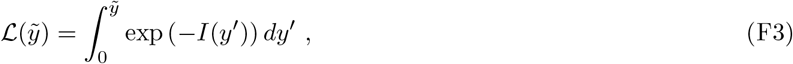

and

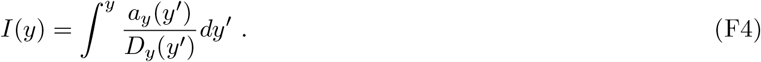

The invasion probability for the first species, *p*_1 fix_, is the fixation probability starting from one individual *n*_1_ = 1, then 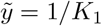. Since 1*/K*_1_ *«* 1 we can try to compute it by expanding in the Taylor series the fixation probability around zero. In this way, we can simplify the integrals (which can be expressed exactly as a combination of hypergeometric functions):

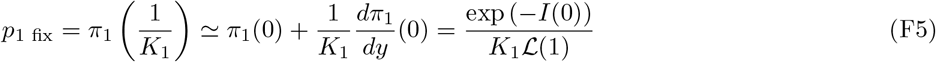

One can obtain also the invasion probability of the second species by noting that the fixation probability reads 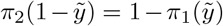. The fixation probability has to be evaluated at 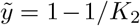, when there is only one individual of the second species and the first one is at carrying capacity. Similarly as before, one can obtain the invasion probability by expanding for 1*/K*_2_ *«* 1:

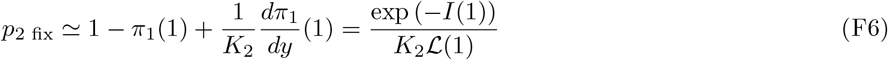

If one considers the ratio between these two probabilities (and neglecting the terms *O*(1*/K*_2_)) the dependency with the integral *ℒ* disappears:

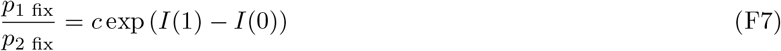

which leads to

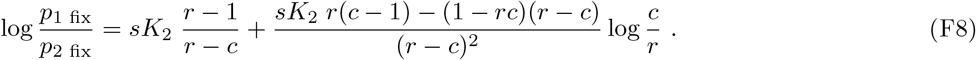

One can notice that in the case *r* = *c* this expression is odd-defined. The case *r* = *c* can be obtained by repeating the full calculation for *r* = *c*. One obtains

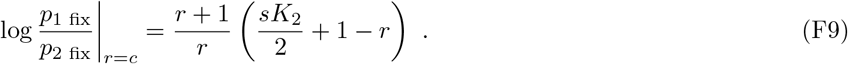

## Appendix G: Simulation of the invasion probability

All the trajectories used in the simulations are generated through a standard Gillespie algorithm for the birth-death process under study.

A straightforward approach can be to generate *R* independent trajectories, where the first strain starts at *n*_1_ = 1, and the abundance of the second strain is sampled from the stationary distribution of the process without type 1: 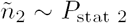. The invasion probability is then just

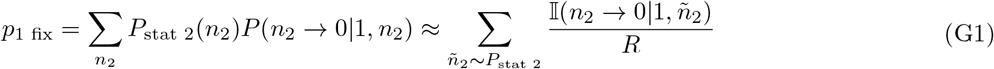

where *P* (*n*_2_ *→* 0|*n*_1_, *n*_2_) represents the probability of the event *n*_2_ *→* 0, i.e. the second population gets extinct, given an initial number of individuals (*n*_1_, *n*_2_). 𝕀 (*n*_2_ *→* 0|*n*_1_, *n*_2_) is the indicator function that is 1 if the event *n*_2_ *→* 0 for a trajectory starting from (*n*_1_, *n*_2_) does occur. However, this approach is very inefficient, since these counts in a simulation are very small (for the parameter setting typically considered, an invasion event is very rare), and a long computational time is needed to have a sufficient amount of samples.

We consider an alternative approach. First, one can observe that, using Bayes rule and the fact that the process is Markovian, the invasion probability can be rewritten by splitting the extinction of the second strain in the probability that the first species reaches a given threshold *n*_1_ = *k* or not:

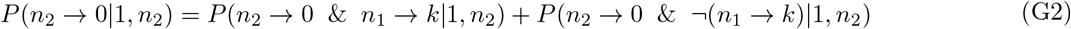

where & is the logical and, and *¬* is the logical negation. The event *n*_1_ *→ k* means that the trajectory has reached (at least once in the past) the value *k*. One can then split the simulation in two parts. One simulates *R* trajectories starting from *n*_1_ = 1 and *n*_2_ sampled from *P*_stat 2_ as before. Then one counts the events of extinction of the second species when the first one does not cross *k*, giving an estimate of the second addendum above. Otherwise, if *n*_1_ reaches *k*, the simulation is stopped, and the number of individuals of the second type is recorded in a set of values 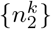. The second part of the simulation consists in generating new *R* trajectories starting from *n*_1_ = *k* and *n*_2_ sampled at random from the list 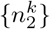. Note that *R* is greater (or equal) than the number of trajectories that in the previous simulation have reached *n*_1_ *→ k*. However, the sampling from 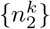 ensures that the statistics of *n*_2_ when *n*_1_ hits *k* is reliable, and, therefore, these new trajectories behave like continuations of the previous ones. Given this new set of *R* trajectories, the count of the events of the second species extinction estimates the first addendum of the equation above. Note that, in this way, when *n*_1_ starts from *k*, the extinction probability is larger, and the counts for its computation increase, leading to better precision. In symbols:

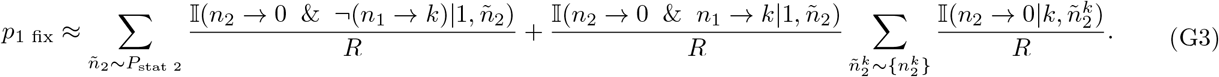

To further improve the statistics, one can add several increasing thresholds (*k*_1_, … *k*_*K*_), and the invasion probability can be obtained with the following iterative formula:

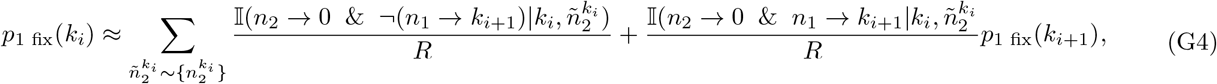

where *p*_1 fix_(*k*_0_) = *p*_1 fix_.

## Appendix H: Evolution in the chemostat model

We consider the chemostat model

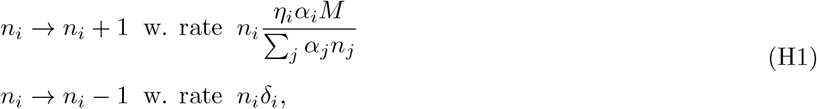

with one constraint to the parameters of the model: *δ*_*i*_*/η*_*i*_ = *α*_*i*_ + *α*_0_, with *α*_0_ *«* 1. The question that we address is which evolutionary outcome one can expect when the parameter *α* can mutate. The answer depends on the details of how the two parameters *δ*_*i*_ and *η*_*i*_ depend on *α*, and, also, if one considers invasions through or without mutations.

Let us first map the chemostat parameters into the general framework. The carrying capacity ratio *c* and the parameter *sK*_2_ read: 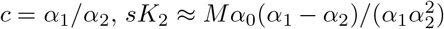. The timescale ratio *r* depends instead on how *δ* and *η* depends on *α*. In particular, we need to separate the cases in which the dependency is contained in the death rate or in the efficiency.

In the main text we consider the case *δ*_*i*_ = *η*_0_(*α*_*i*_ + *α*_0_) and *η*_*i*_ = *η*_0_, which corresponds to, at the leading order in *α*_0_, *r* = *c* = *α*_1_*/α*_2_, and leads to eq. 13

The other case considers a constant death rate *δ*_0_, and *η*_*i*_ = *δ*_0_*/*(*α*_*i*_ *− α*_0_). This leads to *r* = 1 and *c* = *α*_1_*/α*_2_. Using equation 10 we obtain the ratio of the invasion probabilities is larger than one if *α*_2_ *< α*_1_ *< Mα*_0_*/α*_2_, The evolutionary equilibrium situation is obtained when 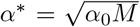, which differs from the constant-efficiency case by a constant factor. A different scenario appears if one instead considers invasions through mutation, as described by eq 11. One can obtain that the invasion is always possible if *α*_1_ *> α*_2_, implying that evolution drives the system to larger and larger values of the intake rate *α*.

### 1. Deriving the distribution of intake rate

In the simulation of figure 4 of the main text, *α* can mutate taking discrete-equispaced values, *{α*_*i*_*}*, with a very small mutation probability that guarantees that the periodic selection regime holds. In this case, one can predict the distribution *p*(*α*) at equilibrium assuming a jump process which satisfies the detailed balance: *p*(*α*_*i*_)*p*(*α*_*i*_ *→ α*_*i*+1_) = *p*(*α*_*i*+1_)*p*(*α*_*i*+1_ *→ α*_*i*_). The transition probabilities are exactly the invasion probabilities through mutations of equation 11:

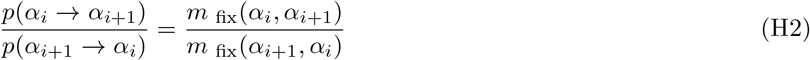

where *m* _fix_(*α*_1_, *α*_2_) is the invasion probability through mutations of a population having intake rate *α*_1_ in a resident population having *α*_2_. Since this ratio is given by equation 11 and 10, one can derive all the ratios between consecutive rates *p*(*α*_*i*_)*/p*(*α*_*i*+1_). All these conditions plus the normalization allow finding all the values of the probability distribution *p*(*α*_*i*_).

